# A unique Cysteine-type protein domain regulates cuticular extracellular matrix assembly in nematodes

**DOI:** 10.64898/2026.02.24.707637

**Authors:** Linxuan Li, Kaiyu Qiu, Hanh Witte, Jörg Martin, Andrei N. Lupas, Ralf J. Sommer

## Abstract

Apical extracellular matrices (aECMs) act as barriers against pathogens and shape tissue organization throughout animals. In nematodes, the cuticle is the aECM performing this function, representing a defining feature of this largest animal phylum. Morphological and molecular evidence supports that worm cuticles extend from the head into the mouth, the latter contributing to the large diversity of nematode feeding structures. However, the biochemical understanding of the compositions of nematode cuticles and head structures is largely limited to collagens. Here, we characterized a recently identified mucin-type protein DPY-6 as a constituent of the cuticle and mouth in the model organisms *Pristionchus pacificus* and *Caenorhabditis elegans*. Utilizing bioinformatic tools, we discover a unique cysteine-consisting motif at the N-terminus of DPY-6 with the amino acid sequence CxCxCxC. We demonstrate through *in vitro* biophysical experiments that this cysteine motif facilitates the intermolecular dimerization of DPY-6 proteins. *In vivo* studies in both species reveal that this motif is involved in the proper localization of the protein in the cuticle, but functions synergistically with other protein domains in a species-specific manner. Given that *dpy-6* transcriptomic expression precedes other cuticle components, we speculate that DPY-6 acts as scaffold molecule for nematode cuticular aECM formation.

## Introduction

Extracellular matrix is defined as a complex heterogenous non-cellular network that is made mostly of macromolecules^1,2^. It has multiple functions across organisms, such as providing physical scaffolds, facilitating cell signaling, and forming barriers^3–5^. One well-studied category are epithelial-covering extracellular matrices, which based on the epithelial surfaces they contact are divided into two groups: apical extracellular matrices (aECMs) and basal extracellular matrices. Basal extracellular matrices attach epithelial basal surfaces, whereas aECM lines with epithelial apical surfaces^6^. As the first line of defense to protect organisms from environmental challenges, e.g. pathogen attack, physical stressors, mechanical injury or ultraviolet radiation, the integuments covering animal surfaces are examples of aECMs serving a barrier function^7,8^. The molecular composition of this type of aECMs varies among animals. In mammals, birds, and reptiles, the skin contains keratin proteins and lipids^9–14^. In contrast, insects and crustaceans deploy chitin within their exoskeletons^15,16^, indicating that aECMs have a more complex and dynamic composition, compared to the conserved basal extracellular matrix^6,17^.

In nematodes, especially in the model organism *Caenorhabditis elegans*, recent progress has identified two separate aECMs: cuticle and pre-cuticle^18^. While the cuticle has long been recognized, only more recent studies identified the pre-cuticle as a separate entity. The pre-cuticle is a transient structure that coats epidermal surfaces preceding the formation of new cuticle. During embryonic morphogenesis and all postembryonic molts, a new pre-cuticle is always formed, and it is endocytosed as the cuticle is built^18,19^. Both structures, the cuticle and the pre-cuticle, share several functional and molecular features with mammalian aECMs, i.e. they contain collagens and other glycoproteins as well^20–23^. Furthermore, traditional morphological studies in nematodes suggested that the cuticle continues into the cheilostome, the most anterior part of the buccal cavity (the mouth of nematodes)^24^. This perspective has recently been supported through findings at the molecular level^25^. Together, the robustness of the cuticle, representing one of the nematode-defining features, and the complicated mouth structures in many nematodes, contribute to the evolutionary and ecological success of worms.

While *C. elegans* and other rhabditid nematodes form a simple mouth structure, which is best described as a tube to ingest bacteria, other nematodes, such as *Pristionchus* and its relatives, develop a more sophisticated buccal cavity (Fig. 1 a-c). *Pristionchus pacificus* has been developed as a second nematode model organism sharing many functional tools with *C. elegans*, but it exhibits unique morphological and behavioral traits that are unknown from *C. elegans* and most other nematodes^26,27^. This includes the development of teeth-like denticles in two different mouth-forms, exemplifying an example of developmental plasticity^26,27^. Depending on environmental factors, *P. pacificus* would form one of two alternative mouth forms, the eurystomatous (Eu, Fig. 1b) or stenostomatous (St, Fig. 1c) form^28^. This developmental process is irreversible. The feeding structures and strategies associated with these buccal cavities are also distinct^28^. *P. pacificus* individuals with a St morph have only one tooth and a narrow buccal cavity. In contrast, the Eu morph has two teeth and a broad buccal cavity. While St animals are strict bacterial feeders, Eu worms are omnivores and can supplement their bacterial diet with predation on other nematodes^28^. This dimorphism of the *P. pacificus* buccal cavity has been intensively studied over the last decade, making this worm a model system for the molecular and genetic analysis of developmental plasticity. Indeed, recent investigations focused on the identification of the regulatory mechanisms and the ecological and evolutionary significance of mouth-form plasticity^29^.

**Figure 1.**
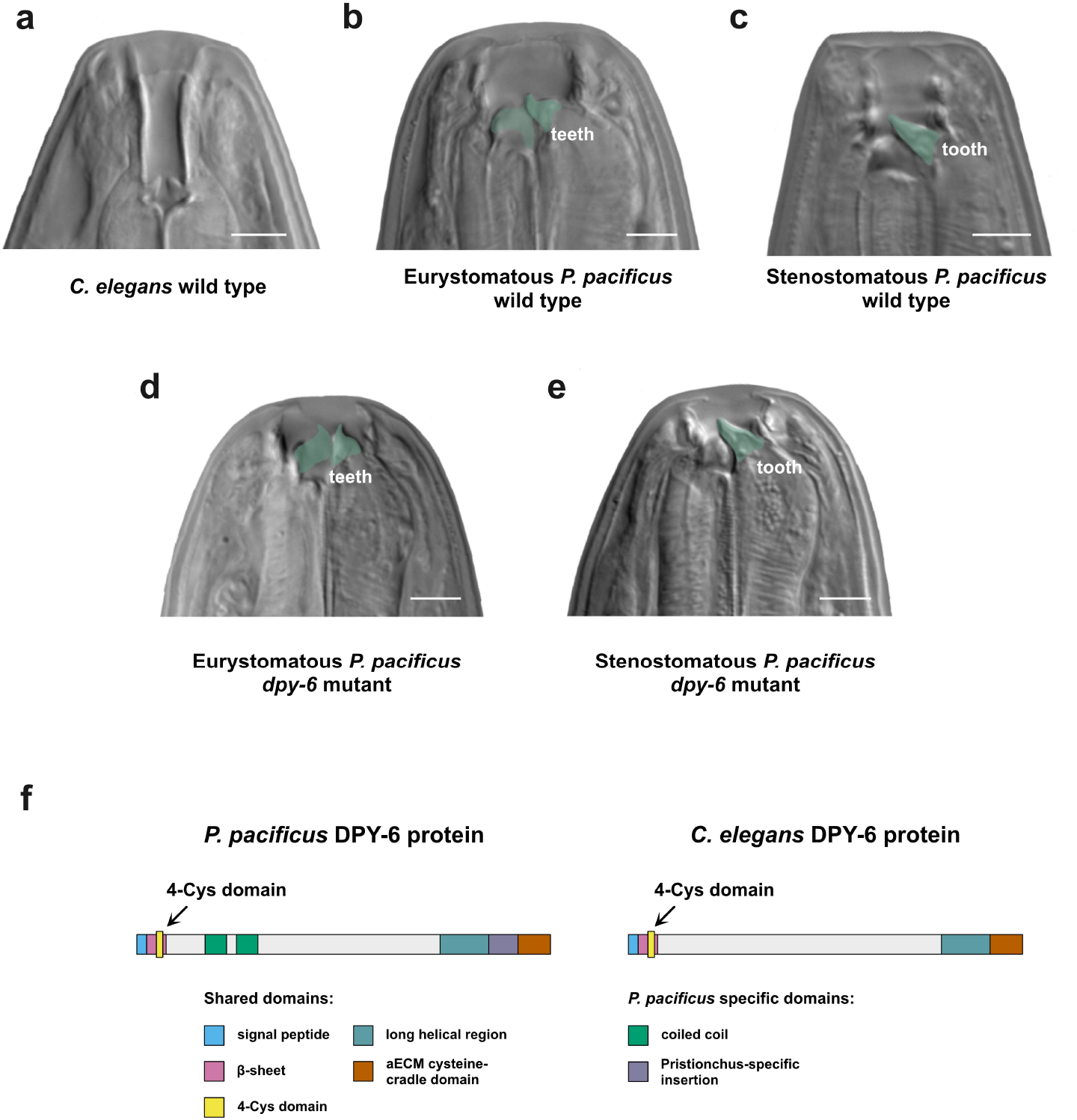
Mouth morphology and DPY-6 protein domains in *C. elegans* and *P. pacificus*. (a) *C. elegans* mouth representing a tube-like structure. (b) Eurystomatous (Eu) mouth form of *P. pacificus*, with two teeth and a wide buccal cavity. (c) Stenostomatous (St) mouth form of *P. pacificus*, with a single tooth and a narrow buccal cavity. (d,e) A *Ppa-dpy-6* mutant worm strain (*tu1768*) has defects in the buccal cavity, with a reduced cheilostome in both the Eu (d) and the St (e) morph. (f) Schematic representation of domain composition of DPY-6 proteins in *P. pacificus* (left) and *C. elegans* (right). Exact amino acid boundaries of each protein domain are represented in Supplementary Data T1. Scale bars for all images are 5μm.

Previous studies have identified various components of the cuticle, including collagens^20,30^, zona pellucida (ZP) domain proteins^31^, chondroitin proteoglycans^32^, and glycoproteins^25,33^. As one type of glycoproteins, mucins are important players of the extracellular matrices in the mammalian respiratory tract and gut^34–36^. In addition, mucins are also found in nematode cuticles. These proteins are rich in serine and threonine residues^37^, which are generally considered as gel-forming components that facilitate the absorption of water and prevent desiccation of the organism^18,22^. *C. elegans* contains more than 20 mucin-encoding genes in its genome^38^. Among them, the *dpy-6* gene is evolutionarily conserved, encoding a mucin-like protein that possesses more than 1200 amino acids and several different functional protein domains. In *C. elegans*, when the *dpy-6* gene is mutated, resulting mutant worms would exhibit a *Dumpy* phenotype with a shorter body size. Compared to other genes causing *Dumpy* phenotype, which normally encode collagens and collagen-modifying enzymes^39^, *dpy-6* is one of only a few non-collagen encoding genes (except for the *C. elegans*-specific dosage compensation genes). Previous literature characterized a novel cysteine-cradle domain (CCD) at the C-terminus of *Cel*-DPY-6 protein^40^. This domain is shared by six genes in *C. elegans*, including *spia-1*^40^. Another recent genetic study identified the orthologous *dpy-6* gene in *P. pacificus*, but *dpy-6* mutants of *P. pacificus* revealed a more complex phenotype^25^. Specifically, *Ppa-dpy-6* mutants are *Dumpy*, similar to *Cel-dpy-6* mutants. However, they also exhibit defects in the formation of the mouth structures, with defects in the anterior part of the buccal cavity, the cheilostome (Fig. 1d and 1e). Note that these findings represent the first genetic and molecular evidence indicating that the cheilostome is continuous with the cuticle and separated from the rest of the buccal cavity^25^.

Here, we combined bioinformatic, biophysical, and genetic analyses to illustrate the function of a unique 4-Cys domain near the N-terminus of DPY-6 proteins. We demonstrate *in vitro* that its functions include intermolecular dimerization. Furthermore, *in vivo* experiments in *C. elegans* and *P. pacificus* revealed that the disruption of this 4-Cys domain led to assembly defects of the cuticle aECM. Thus, our findings characterize functions of a novel protein domain that we named “disulfide staple domain”, indicating its important roles for the organization of the nematode cuticle.

## Results

### A novel protein domain containing an alternating pattern of four Cysteines is restricted to nematodes

With multiple different protein domains, the *P. pacificus* and *C. elegans* mucin-type DPY-6 proteins are extremely large. Figure 1f provides a schematic representation of the predicted DPY-6 protein domains from *P. pacificus* and *C. elegans*. At the N-terminus, both DPY-6 proteins have a signal peptide, followed by a conserved four-cysteine motif residing in a β-sheet (Fig. 1f). After this unique structure, the *Ppa*-DPY-6 contains two predicted coiled-coil domains (H1 and H2) that are not conserved in *Cel*-DPY-6. A large unstructured region follows behind, and then both proteins have a long predicted helical region. In *P. pacifics*, this domain is accompanied by a *Pristionchus*-specific insertion. At the C-terminus of both DPY-6 proteins is a novel alpha-helix bundle, recently named “aECM cysteine-cradle domain” (aCCD) after its original characterization in *Cel-* SPIA-1^40^. Thus, the *Cel*-DPY-6 protein shares most of its domains with *Ppa*-DPY-6, except for two coiled-coil domains and the *Pristionchus*-specific insertion. However, little is known about the biochemical and biophysical functions of DPY-6 proteins in the formation of the cuticle and the buccal cavity.

We focused on the 4-Cys motif of DPY-6 proteins, which we initially referred to as the “4-Cys domain” (Fig. 1f). The arrangement of these four cysteine residues is conspicuous, as they are separated from each other by one single amino acid, resulting in a “CxCxCxC” pattern (Fig. 2a). Because of the alternating pattern in which amino-acid side chains project from a β-sheet, this ordering causes the cysteine sulfhydryl groups to all point in the same direction. Therefore, these cysteines cannot form disulfide bonds intramolecularly. Sequence similarity searches showed us that this domain is conserved across nematodes and not found outside this phylum. Thus, nematode DPY-6 proteins contain a novel domain of unknown function with an unusual arrangement of four alternating cysteine residues.

**Figure 2.**
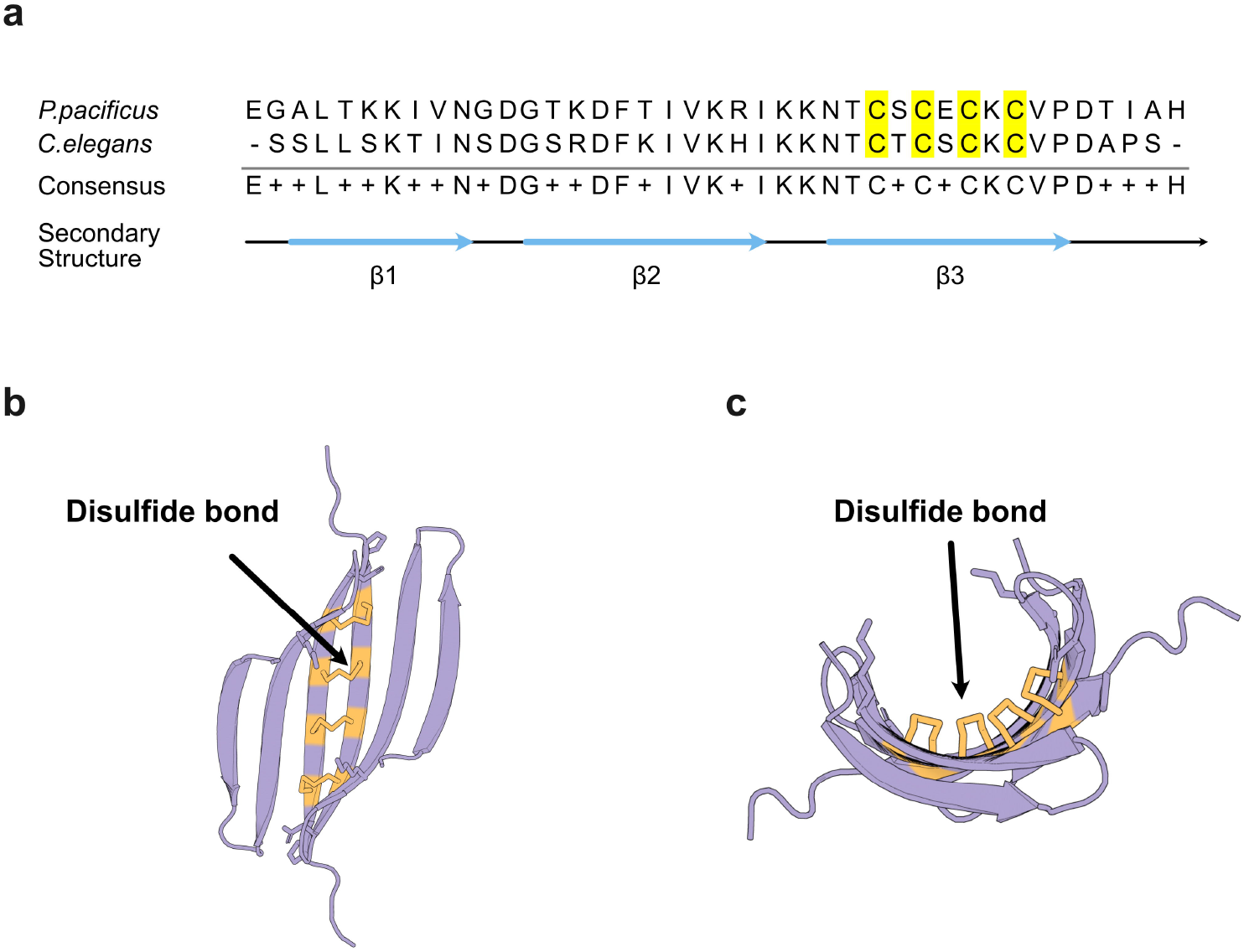
Amino acid sequences and AlphaFold folding prediction of the “4-Cys domain” (disulfide staple domain). (a) Amino acids sequence alignments of β-sheet in DPY-6 proteins from *P. pacificus* and *C. elegans*. In this β-sheet, four conserved alternating cysteines constitute the “4-Cys domain”. (b) AlphaFold predictions of the dimerized status of the “4-Cys domain” from *P. pacificus*. This dimerization is held by the disulfide bonds formed intermolecularly between two “4-Cys domains”. (c) Same dimer viewed from a lateral perspective.

### AlphaFold predicts dimerization of two antiparallel “4-Cysteine domains”

Using AlphaFold predictions, we investigated the potential oligomerization status of the 4-Cys domain. Indeed, AlphaFold supports a potential dimerization of the 4-Cys domain, with two monomers running antiparallel to each other (Fig. 2b). In this scenario, the cysteine residues would be able to form disulfide bonds intermolecularly to lock monomers together. Furthermore, this model also implicates that the dimerized motif would form a cradle-like fold (Fig. 2c), given the curvature of the β-sheet. Note that this AlphaFold prediction possesses the predicted template modelling (pTM) score as 0.61 and the interface predicted template modelling (ipTM) score as 0.58 (Supplementary Data S-Figure 1), indicating need for experimental verification^41^.

### Biophysical evidence supports the dimerization of the 4-Cysteine domain

We used several biophysical approaches to test the AlphaFold predictions. First, we expressed recombinant proteins with the amino acid sequence of the β-sheet containing the 4-Cys motif from *C. elegans* and *P. pacificus* in SHuffle cells, an engineered *E. coli* BL21 derivative^42^. This cell expression system provides a suitably oxidizing cytosolic environment for the formation of disulfide bonds. The recombinant proteins were named “Cel_4cys” and “Ppa_4cys”, respectively (Fig. 3a). After purification, we performed circular dichroism (CD) measurements to establish the secondary structure of the recombinant proteins. As shown in Figures 3b and 3c, both Cel_4cys and Ppa_4cys exhibited β-sheet CD patterns. This finding not only supported the AlphaFold predictions, but also indicated that the recombinant proteins produced in SHuffle cell were properly folded.

**Figure 3.**
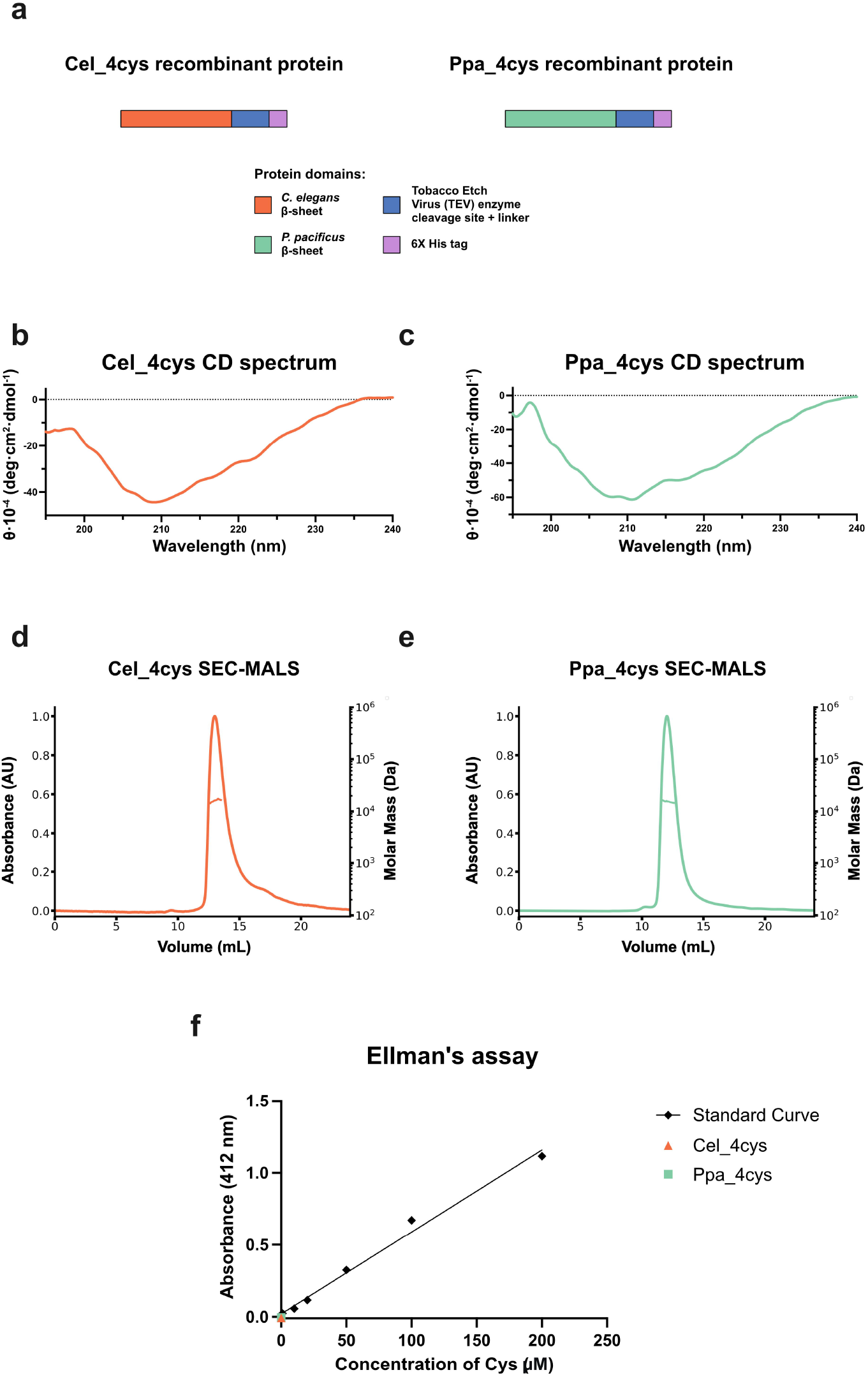
*In vitro* experiments related to recombinant proteins Cel_4cys and Ppa_4cys. (a) Domain compositions of Cel_4cys and Ppa_4cys recombinant proteins. (b) CD spectrum results of Cel_4cys, showing a β-sheet pattern. (c) CD spectrum results of Ppa_4cys, showing a β-sheet pattern. (d) SEC-MALS data of Cel_4cys with a single peak corresponding to 16.2±0.7kDa, implying dimerization of Cel_4cys. (e) SEC-MALS data of Ppa_4cys with a peak corresponding to 16.0±0.6kDa, implying dimerization. (f) Ellman’s assay of Cel_4cys and Ppa_4cys. Both proteins have absorbances close to the one of 0μM in the standard curve, suggesting that there is nearly no free cysteines.

Next, we performed <u>S</u>ize-<u>e</u>xclusion <u>c</u>hromatography-<u>M</u>ulti<u>a</u>ngle <u>L</u>ight <u>S</u>cattering (SEC-MALS) experiments. SEC-MALS results with the Cel_4cys protein displayed one single peak, which corresponded to a protein size of 16.2±0.7kDa (Fig. 3d). This experimental outcome suggests that the recombinant protein exists in solution as one single stable form. Given that the monomer of the Cel_4cys protein has a size of 6.6kDa, the SEC-MALS result implied that the Cel_4cys proteins formed dimers in solution. The Ppa_4cys protein revealed a similar pattern with a single peak of 16.0±0.6kDa (Fig. 3e). Together, these results support the idea that the nematode-specific 4-Cys domain undergoes dimerization, as predicted by AlphaFold.

We also employed Ellman’s assay to determine if the dimerization process is facilitated by the disulfide bonds formed between the cysteine residues. This assay utilizes 5,5’-dithio-bis-(2-nitrobenzoic acid), known as DTNB, to probe for any free sulfhydryl groups, such as in cystine side chains. DTNB would interact with the free sulfhydryl groups and produce a yellowish compound, 2-nitro-5-thiobenzoic acid (TNB). Through absorbance measurements at 412nm, we found that 40μM recombinant protein samples of Cel_4cys and Ppa_4cys both displayed a relatively low absorbance, similar to the 0μM standard curve control (Fig. 3f). These observations indicate that there were nearly no free cysteine residues within the protein samples. Thus, almost all cysteine residues form disulfide bonds, in agreement with the AlphaFold model. Given that the four cysteine residues function as staples to lock two monomers together, we finally named this domain “disulfide staple domain”.

### An in-frame deletion of the disulfide staple domain disrupts *C. elegans* DPY-6 localization and leads to a *Dumpy* phenotype

We further studied the DPY-6 proteins *in vivo*, making use of the genetic engineering tools available in *C. elegans* and *P. pacificus*. To better track the localization of the DPY-6 protein, we employed ALFA-tagging, a novel protein tag that can be reliably applied in nematodes for either immunostaining or pulldown experiments^43^. We added two copies of the ALFA-tag in the middle of the *Cel*-DPY-6 protein, right after the disulfide staple domain (Fig. 4a). Adding the ALFA-tag did not change the morphology of *C. elegans* worms (Fig. 4c and 4d). Moreover, immunostaining displayed that the *Cel*-DPY-6 protein is localized to the furrows of the cuticle in L4 larvae, exhibiting a parallel stripe pattern (Fig. 4e and 4f).

**Figure 4.**
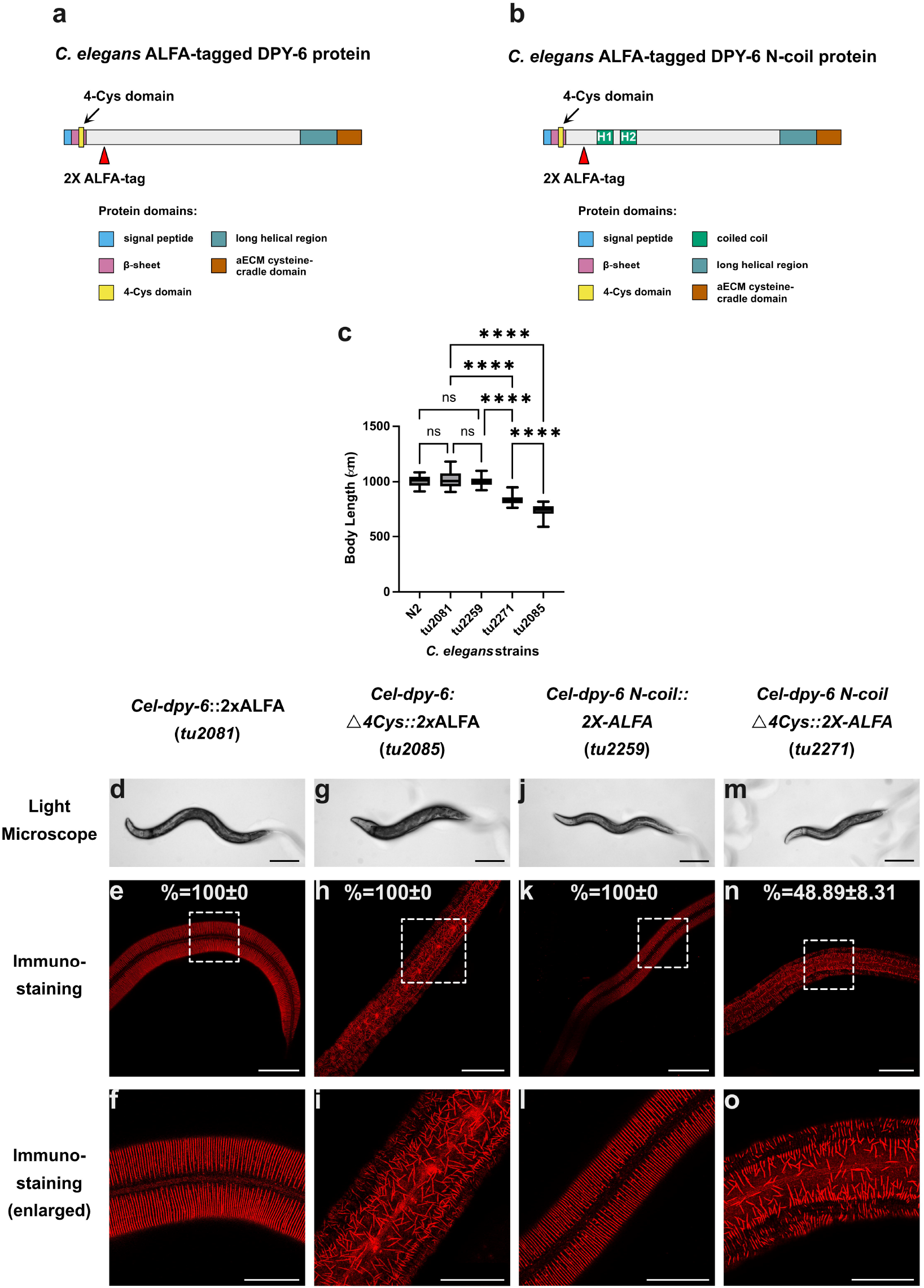
Schematic images for CRISPR engineered proteins, worm body lengths and DPY-6 localization of wild type and *dpy-6* mutant versions in *C. elegans*. (a) Schematic representation illustrating the location of ALFA-tag in the *C. elegans* DPY-6 protein. (b) Schematic representation of a hybrid DPY-6 protein after the insertion of the *P. pacificus* N-terminal coiled-coil domains (H1 and H2) into the *C. elegans dpy-6* locus. (c) Body lengths of wild type and various *C. elegans* mutant strains (One-way ANOVA, **** p<0.0001; n.s., not significant; n=50). (d-o) Light microscopic images (d, g, j, m) and DPY-6 cuticle localization (e, f, h, i, k, l, n, o) of wild type (d-f) and various mutant forms (g-o). Immunostaining was performed using confocal microscopy. Enlargements in f, i, l, and o show areas highlighted by white-dotted squares. The numbers in e, h, k, and n represent the mean value ± the standard deviation of the DPY-6 staining patterns’ percentages in the corresponding stained worm strains. Scale bars in d, g, j, and m are 200μm; in e, h, k, and n are 50μm; in f, i, l and o are 25μm.

We then utilized the *Cel-dpy-6*::2xALFA strain *tu2081* to study the function of the disulfide staple domain *in vivo*. For that, we in-frame deleted the whole β-sheet structure bearing the disulfide staple domain by using CRISPR-Cas9 engineering. The in-frame deletion allows us to specifically remove the target structure, whereas all other protein domains remain intact. We discovered that *Cel-dpy-6:*^△^*4Cys*::2xALFA(*tu2085*) mutant worms displayed a strong *Dumpy* phenotype (Fig. 4c and 4g). Strikingly, immunostaining of these mutant worms revealed that the stripe pattern was completely destroyed (Fig. 4h and 4i). Instead, *Cel*-DPY-6 protein was found to be organized in random short filaments in the cuticle. These findings demonstrate that the disruption of the disulfide staple domain of *Cel-*DPY-6 prevents proper localization of the protein, leading to a *Dumpy* phenotype.

### *Ppa*-DPY-6 shows different extracellular matrix pattern and in-frame deletion of the disulfide staple domain has no phenotype

We conducted similar *in vivo* experiments in *P. pacificus* (Fig. 5). The addition of the ALFA-tag in *P. pacificus* did not influence the body shape of the worm (Fig. 5 a-c). As expected, it also revealed the localization of *Ppa*-DPY-6 protein to the cuticle in J4 larvae. Similar to *C. elegans*, the DPY-6 protein is localized in the furrows of the cuticle (Fig. 5d and 5e). However, in contrast to *C. elegans*, the localization pattern of *Ppa*-DPY-6 is not in parallel stripes, but instead exhibits a pattern of dashed dots assembling into longitudinal lines (Fig. 5d and 5e). This is because *P. pacificus* and *C. elegans* have distinct types of cuticle furrow patterns. While *C. elegans* furrows are circumferential, furrows of *P. pacificus* are evenly distributed in a longitudinal orientation along the body length^44^. Notably, these longitudinal lines are characteristics of the cuticle in the majority of *Pristionchus* species, and they are readily visible by light and electron microscopy. Taken together, cuticle furrows differ distinctively between *C. elegans* and *P. pacificus*, which is also represented by the localization patterns of DPY-6 proteins in the two species.

**Figure 5.**
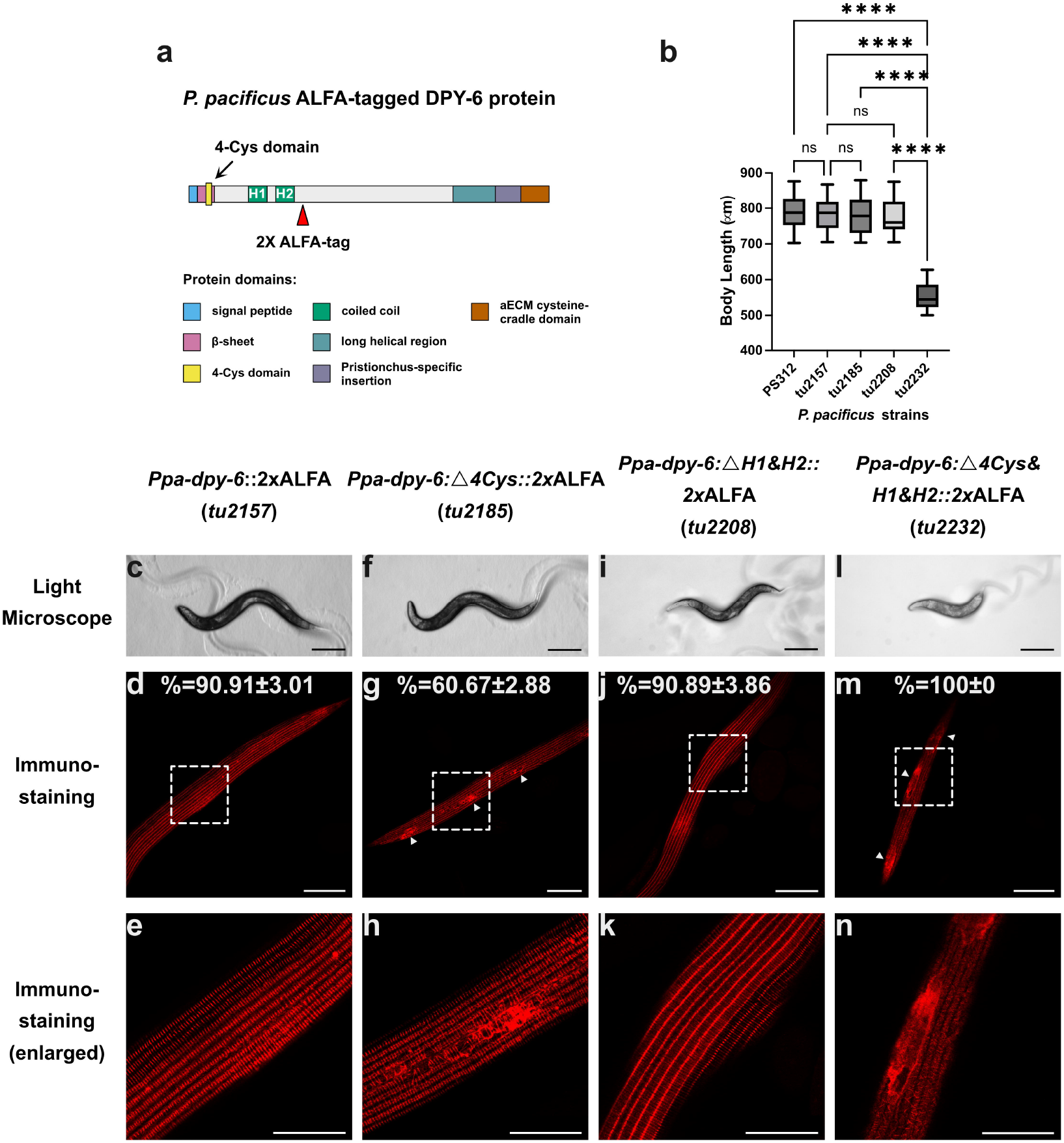
Schematic images for CRISPR engineered proteins, worm body lengths and DPY-6 localization of wild type and *dpy-6* mutant versions in *P. pacificus*. (a) Schematic representation illustrating the locations of ALFA-tag in the *P. pacificus* DPY-6 protein. (b) Body lengths of wild type and various *P. pacificus* mutant strains (One-way ANOVA, **** p<0.0001; n.s., not significant; n=50). Light microscopic images (c, f, i, l) and DPY-6 cuticle localization (d, e, g, h, j, k, m, n) of wild type (c-e) and various mutant forms (f-n). Immunostaining was performed using confocal microscopy. Enlargements in e, h, k, and n show areas highlighted by white-dotted squares. The numbers in d, g, j, and m represent the mean value ± the standard deviation of the DPY-6 staining patterns’ percentages in the corresponding stained worm strains. Scale bars in c, f, i, and l are 200μm; in d, g, j and m are 50μm; in e, h, k and n are 25μm.

Afterwards, we also in-frame deleted the disulfide staple domain in the ALFA-tagged *Ppa*-DPY-6 protein. Surprisingly, we discovered that the removal of the disulfide staple domain did not cause a *Dumpy* phenotype similar to *Cel-dpy-6* mutants (Fig. 5b and 5f). Moreover, immunostaining of these *P. pacificus* mutants showed that the localization of the *Ppa*-DPY-6 protein was only weakly affected (Fig. 5g and 5h). Unlike the *Cel*-DPY-6 mutant protein, which formed random protein filaments all over the cuticle, most of the *Ppa*-DPY-6 mutant protein was still observed at the correct location (Fig. 5g and 5h). We hypothesized that this partially correct localization of *Ppa-*DPY-6 facilitated the maintenance of the worms’ body shape and, thus, the absence of a *Dumpy* phenotype. Note that despite the fact that most of *Ppa*-DPY-6 protein was seen in the furrows, we observed sporadic random protein aggregations on the cuticle in about 60.67± 2.88% of *P. pacificus* mutant animals (Fig. 5g, white arrows). In comparison, this aggregation was only seen in about 9.09±3.01% of full-length ALFA-tagged DPY-6 *P. pacificus* worms. These experimental outcomes imply that the disulfide staple domain in *P. pacificus* either functions together with other domains to control proper localization of the DPY-6 protein, or, alternatively, has no role similar to the one observed in *Cel-*DPY-6.

### The *Pristionchus* disulfide staple domain acts in synergy with two coiled-coil domains to regulate extracellular matrix assembly

The mechanisms underlying these phenotypic differences between *C. elegans* and *P. pacificus* mutants could be due to the presence of the H1 and H2 coiled-coil domains, which are only found in *Ppa-*DPY-6 (Fig. 1f). These two coiled-coil structures could potentially assist dimerization and indeed, bioinformatic predictions support the dimerization of these coiled-coil domains^45^. Therefore, one hypothesis would be that the coiled-coil domains might function redundantly with the disulfide staple domain to guarantee the dimerization of the *Ppa*-DPY-6 protein.

To test this hypothesis *in vivo*, we firstly in-frame deleted both H1 and H2 structures by CRISPR-Cas9 engineering in the DPY-6 ALFA-tag background. Resulting *Ppa-dpy-6*^△^*H1&H2*(*tu2208*) mutant worms did not exhibit any phenotypic change with respect to body shape (Fig. 5b and 5i). Similarly, immunostaining also did not show any obvious changes in the localization of the *Ppa*-DPY-6 protein in the cuticle (Fig. 5j and 5k). In contrast, when we introduced the in-frame deletion of the disulfide staple domain into *Ppa*-*dpy-6*^△^ *H1&H2* mutant worms, we found that the resulting mutant worms (*tu2232*) displayed a strong *Dumpy* phenotype (Fig. 5b and 5l). In addition, immunostaining indicated that the localization of the mutated *Ppa*-DPY-6 protein was severely disrupted (Fig. 5m and 5n). Specifically, the mutated *Ppa*-DPY-6 protein was observed either as a smear or in an aggregated state (Fig. 5m, white arrows). These experiments indicate that the correct localization and hence the proper function of the DPY-6 protein in *P. pacificus* requires the synergistic interaction of the disulfide staple domain with the coiled-coil domains.

Finally, we wanted to investigate if the addition of the *P. pacificus* H1 and H2 coiled-coil domains could affect the *Cel-*DPY-6 protein. To examine this, we added these two domains into *Cel*-DPY-6 (Fig. 4b). When inserting these domains into an otherwise wild type *Cel-*DPY-6 protein, we noticed no obvious change (Fig. 4c, 4j-l). However, when we introduced the in-frame deletion of the disulfide staple domain into this background, the new mutant line *Cel-dpy-6 N-coil*^△^*4Cys*::2xALFA(*tu2271*) still showed a *Dumpy* phenotype (Fig. 4c and 4m). Note that the *Dumpy* phenotype is not as strong as that of other alleles (Fig. 4c and 4m). Immunostaining experiments also showed that even if the majority of mutant worms still possess randomly organized short filaments, in about 48.89± 8.31% of the animals the wild-type parallel stripe patterns of DPY-6 protein was seen to some extent (Fig. 4n and 4o). These observations are different from the corresponding manipulations in *P. pacificus*, implying that the addition of the coiled-coil domains could not fully rescue the *Dumpy* phenotype in *C. elegans*. In summary, these results suggest that the *P. pacificus* coiled-coil domains of DPY-6 have changed the functional properties of the otherwise conserved DPY-6 protein.

## Discussion

In this study, we identified a novel domain in the nematode mucin-type protein DPY-6 named “disulfide staple domain”, and characterized its functional properties utilizing biophysical and genetic experiments. Recent work in *C. elegans* has indicated that the nematode aECM has to be divided into pre-cuticle and cuticle, which play important roles for the development, physiology and behavior of the organism^18^. The dynamic features of both of these structures represent an excellent model for mammalian aECM^18^. However, the highly-dynamic assembly and disassembly processes of nematode aECMs have to be seen in the context of molting. Like other ecdysozoans, nematodes have to shed their cuticles multiple times during development. Before reaching adulthood, all nematodes need to pass through four juvenile stages J1-J4 (named L1-L4 in *C. elegans*), with each stage separated by a molt. During molting, the old cuticle is replaced by a new cuticle, requiring massive protein production in the hypodermis. Much of this protein production occurs during a quiescent phase preceding molting, called lethargus^46^. While this process is long known, it is only recently that a separate aECM structure, the so-called pre-cuticle, was discovered in *C. elegans* to preceded to the new cuticle in each molt^18^.

Many studies in the past focused on the collagen-encoding genes, showing that different collagens have different expression profiles, and immunolocalization studies further support sub-functionalizations of the investigated collagens^18,47^. On the contrary, work on non-collagen proteins in the aECM is limited. Our research presented here highlights critical functions of a non-collagen protein in aECM formation in *P. pacificus* and *C. elegans*. Using bioinformatic, biophysical and molecular genetic tools, we obtained four novel insights through our work, which are discussed below.

First, our findings indicate a fundamental role of the newly identified CxCxCxC domain of DPY-6 (disulfide staple domain) for the assembly of nematode cuticular aECMs. Various biophysical experiments support intermolecular disulfide bond formation between two antiparallel DPY-6 proteins, which is in line with the AlphaFold prediction (Figures 2 and 3). In contrast, our CRISPR-engineered *Ppa-dpy-6* mutants lacking the disulfide staple domain alone do not show any defects in buccal cavity and teeth formation in *P. pacificus* (Supplementary Data S-Figure 2). As *Ppa-dpy-6* knockout mutants have a strong phenotype in the mouth (Fig. 1d and 1e), these findings might indicate a role of other DPY-6 protein domains in cheilostome formation. Therefore, the absence of a mouth phenotype after the in-frame deletion of the disulfide staple domain was surprising and suggests further intramolecular redundancies. Ongoing studies of other *Ppa-*DPY-6 domains might provide novel insights in this context.

Second, our work provides novel insights into the rapid diversification of nematode aECM proteins. While many aECM components turn over rapidly at the level of individual genes^48,49^, i.e. collagens, the *dpy-6* locus is conserved throughout nematodes as a single copy gene. However, the comparison between the *C. elegans* and *P. pacificus* DPY-6 proteins exhibits different domain compositions, i.e., the emergence of a coiled-coil domain and a *Pristionchus*-specific insertion in *Ppa*-DPY-6. Genetic investigations through in-frame deletions described in this study support the functional importance of such novel domains in *P. pacificus*. Thus, the DPY-6 protein and its interaction partners might be rapidly evolving.

Third, given the structural properties of DPY-6 and the observed mutant phenotypes in both species, we hypothesized that DPY-6 serves as a scaffolding protein, on which other proteins, such as collagens, can start to form the complex aECM structure. This hypothesis is supported by transcriptomics indicating that *Cel-dpy-6* expression precedes other genes encoding constituents of the pre-cuticle and cuticle in each molting cycle^40,50^. Similarly, high-resolution developmental transcriptomics in *P. pacificus* also found that *Ppa-dpy-6* expression precedes other aECM components^51^. The evolutionary conservation of the disulfide staple domain throughout nematodes further supports its importance for proper cuticular aECM assembly. At the same time, the largely unstructured regions of DPY-6 protein allow extensive glycosylations, which then lead to the gel-forming properties of mucin-type proteins. This could also be critical for the scaffolding function of DPY-6. Additionally, recent experiments in *C. elegans* are also consistent with a scaffolding function of DPY-6^52^. However, additional biophysical and genetic studies are necessary to directly test this hypothesis.

Finally, it will be interesting to investigate the potential binding partners of the dimerized disulfide staple domain. The dimerized motif forms a cradle-like fold (Fig. 2c), given the curvature of the β-sheet. This dimerized structure could potentially bind to molecules possessing linear structures. Besides proteins, recent investigations also discovered that chitin is one of the major building blocks of both forms of teeth in *P. pacificus*^53^. When the chitin synthase-2 gene *Ppa-chs-2* was knocked out, chitin biosynthesis defective worms became teethless and revealed defects in the predatory behaviors. Nevertheless, *Ppa-chs-2* mutants can be fed normally on a bacterial diet. Considering that chitin is a linear polysaccharide and exists in parts at the same location as the *Ppa*-DPY-6 protein^40^, we would hypothesize that chitin might be a potential binding partner for DPY-6 proteins through the disulfide staple domain. Further experiments will be necessary to test this hypothesis, such as Isothermal Titration Calorimetry or Nuclear Magnetic Resonance (NMR) spectroscopy. Note that previous studies only detected chitin in the teeth and eggshell of *P. pacificus*^53^ and in the eggshell and pharynx of *C. elegans*^54^, not in the cuticle of either species. Therefore, it is well possible that other non-chitin interacting partners of DPY-6 exist in the cuticle.

In conclusion, the biophysical and genetic analysis of a unique cysteine-composing protein domain specific to nematodes has revealed novel insight into the formation and regulation of aECM assembly. Moreover, this finding further emphasizes the suitability of nematodes as a model system in aECM-related work, owing to their optical transparency and the relative ease of generating CRISPR-engineered mutants with various targeted protein domain modifications.

## Author Contributions

Conceptualization: R.S. and A.L.; Bioinformatics: A.L. and K.Q.; *in vitro* experiments: L.L. and J.M.; *in vivo* experiments: L.L. and H.W.; Writing: L.L., R.S., and A.L.

## Acknowledgement

We would like to thank Dr. Nathalie Pujol for constructive suggestions about the manuscript. L.L. would like to express his thanks to Dr. Cátia Igreja, Radhika Sharma and Dr. Yimin Hu for detailed project discussions and skill teaching, Aurora Panzera for the help of imaging acquiring, as well as Anja Rau, Eva Hertle, Astrid Ursinus and Dr. Birte Hernandez Alvarez for technical support. Finally, L.L. would like to give his highest gratitude to Mr. Zheng’ao Li, who led L.L. into the world of science, and Ms. Fengying Li, who accompanied L.L. through his way to the dream.

## Methods

### *P. pacificus* and *C. elegans* maintenance

All *P. pacificus* and *C. elegans* strains were cultured under standard laboratory conditions using nematode growth medium (NGM) agar plates with *Escherichia coli* OP50 as food source as previously described^39^. Detailed information about all nematode strains can be found in the Supplementary Data T2 and T3.

### CRISPR-Cas9 genetic engineering

Transgenic worms were generated using CRISPR-Cas9 genetic engineering as previously described^55^. In brief, gene-specific crRNAs were designed to target 20 bp upstream of the protospacer adjacent motifs (PAMs). All chemicals were purchased from Integrated DNA Technologies. From 100μM stocks of the crRNA and the tracrRNA (catalog# 1072534), 10μL solutions were obtained and mixed together, incubated at 95 °C for 5 min for denaturation and then cooled down to room temperature for annealing. Afterwards, Cas9 protein (catalog# 1081058) was added to the mixture and allowed to incubate at the room temperature for another 5 min. TE buffer was used to dilute the solution so that the final concentration of 18.1μM RNA hybrid and 2.5μM Cas9 protein were reached. To achieve specific site-directed mutation, repair templates in the form of ssDNA were included at a final concentration of 4 µM. Repair templates were designed to contain the desired modifications, flanked by at least 35nt homology arms at both sides of the edited sequence. When necessary, the PAM motif or the gRNA binding sequence was changed to include silent mutations, so that the edited allele was not re-cut. For the addition of the ALFA-tag, the repair templates included two copies of the nucleotide sequences encoding the ALFA-tag, which were codon-optimized for *P. pacificus* and *C. elegans*, respectively. Moreover, the sequences of these two copies of the ALFA were designed with different codons for the same amino acid to minimize sequence identity. As co-injection marker, we used a plasmid containing the *Ppa-eft-3* promoter with a modified TurboRFP sequence^56^. The final solution was injected into the germline of *P. pacificus* or *C. elegans* using a Zeiss Axiovert microscope (Zeiss, Germany) and an Eppendorf FemtoJet injector (Eppendorf AG., Hamburg, Germany). Injected P0 worms were allowed to lay eggs for 12-16h post injection, then P0 worms were removed and F1 eggs were recovered. After 4 days, plates containing F1 worms were screened for fluorescent worms indicating transgenic animals. From plates containing fluorescent F1 worms, 12-16 progenies were isolated and singled out. After the F1 animals laid eggs, they were lysed and the genomic region of interest was sequenced by Sanger sequencing (GENEWIZ Germany GmbH)^57^. Worms with desired mutations were further studied to obtain homozygous mutant worms.

### Immunostaining Experiments

Immunostaining experiments with ALFA-tagged worms were conducted as previously described^43^. In brief, worms were collected from NGM plates with M9 medium and washed until bacteria were gone. Fixation was performed with 4% para-formaldehyde, 1xPBS (500 µL) in 1.5 mL Eppendorf tubes overnight on a tube rotator. Samples were washed three times using a washing buffer (0.5% Triton X-100, 1x PBS) and then incubated overnight with 300rpm shaking in 500 µL of buffer containing 5% β-mercaptoethanol, 1% Triton X-100 and 0.1 M Tris-HCl pH 7.4. This step is necessary to reduce disulfide bonds. The next day, samples were washed twice with 1% Triton X-100, 0.1 M Tris-HCl pH 7.4 buffer and once with 1 mM CaCl_2_, 1% Triton X-100, 0.1 M Tris pH 7.4 buffer. Worms were incubated at 37 °C for 2–3h with 1 mM CaCl_2_, 1% Triton X-100, 0.1 M Tris pH 7.4 buffer. Washing steps with washing buffer were performed three times, followed by incubation at room temperature for 1h with blocking buffer (1% BSA, 0.5% Triton X-100, 1x PBS). After two more washing steps, samples were stained with the primary antibody (FluoTag-X2 anti-ALFA-AbberiorStar580, NanoTag Biotechnologies, Göttingen, Germany, 1:100) in 50 µL of blocking solution, overnight at 4 °C in a tube rotator. The next day, animals were washed again with washing buffer three times. Animals were resuspended in VectaShield mounting medium (Vector Laboratories, USA) containing 1 μg/mL 4⍰,6-diamidino-2-phenylindole (DAPI; Molecular Probes, Thermo Fisher Scientific) and mounted on freshly prepared 5% Noble agar pads. A Leica TCS SP8 microscope was used to acquire corresponding pictures, and Fiji (Image J, NIH, Bethesda, MD, USA) was used for the following analysis. For phenotype counting, at least 30 stained worms were included for each replicate, three replicates were conducted in total.

### Worm size measurements

Young adult hermaphrodites from each *C. elegans* and *P. pacificus* strains were moved to NGM plates without bacteria. Bright-field images were taken at ×10 magnification using the Axio Zoom V16 microscope. 50 worms were measured for each strain. Measurement was done on the software ZEN 3.3.

### Bacterial strains and cultivation

Molecular cloning was performed using NEB® 5-alpha competent *E. coli* cells (Cat. C2987H, New England Biolabs, Ipswich, Massachusetts, United States). For protein expression, SHuffle® T7 Express competent *E. coli* (Cat. C3029J, New England Biolabs, Ipswich, Massachusetts, United States) was chosen. Bacteria were grown in Luria-Bertani (LB) media supplemented with appropriate antibiotics.

### Cloning and plasmids

Both sequences of the “4-Cys domain” (disulfide staple domain) in *P. pacificus* and *C. elegans* DPY-6 proteins were codon-optimized and then synthesized either by Integrated DNA Technologies, Coralville, Iowa, United States, or by ThermoFisher Scientific. Both proteins were expressed, fused to a TEV enzyme cleavage site sequence and an C-terminal (Histidine)_6_ tag, using the vector pET-28a(+) plasmid.

### Protein expression and purification

Proteins were expressed in SHuffle® T7 Express competent *E. coli*. LB media supplemented with kanamycin was used for bacterial cultures. Cultures were incubated at 30 °C until an optical density of OD600 = 0.6 was reached. Isopropyl-β-d-thiogalactoside (IPTG) was added to a final concentration of 1 mM to induce protein expression. After that, bacterial cultures were agitated at 16°C for 40 h for higher yield. Cells were pelleted and resuspended in lysis buffer (50 mM Tris, pH 8.0, 50 mM NaCl, and 5 mM MgCl_2_, protease inhibitor mix (cOmplete™, EDTA-free Protease Inhibitor Cocktail, Roche), DNase I (AppliChem GmbH). French press was utilized to lyse cells, and cell debris and insoluble material were pelleted by centrifugation at 20,000 rpm for 45 min at 8 °C. The supernatant was filtered through a 0.45 μm filter and a 0.22 μm filter. Afterwards, the solution was applied to a 5 mL HisTrap column (Cytiva). Bound proteins were eluted using a linear imidazole gradient, ranging from 10–600 mM, in the background buffer (50mM Tris, pH 7.0, 300mM NaCl). The eluted protein was cleaved with TEV protease to remove the His-tag. After overnight cleavage, a second purification step was conducted using a HisTrap column to separate the protein from the cleaved tag. The whole step was performed using a buffer containing 50mM Tris, pH 7.0, 100 mM NaCl. Following this step, proteins were purified to homogeneity by utilizing gel filtration chromatography on a Superdex 75 column (Cytiva), which was equilibrated with SEC buffer (50mM Tris, pH 7.0, 100 mM NaCl). Protein purity was confirmed by SDS-PAGE (mPAGE 4–12% Bis-Tris Precast Gel, Millipore) and protein concentration was determined spectrophotometrically.

### Circular dichroism (CD) spectroscopy

Samples were dialyzed against the CD buffer (10mM potassium phosphate buffer, pH 7.0, 50mM Na_2_SO_4_), and diluted to a final concentration of 0.11mg/ml (Cel_4cys) and 0.08mg/ml (Ppa_4cys), respectively. CD spectra were recorded using a Jasco J-810 spectrometer (JASCO). The cuvette used has a 1 mm path length. The wavelength range is 195–240 nm, and 100 nm/min is chosen as reading speed. A total of 10 single spectra were recorded and averaged. To analyze data, we subtracted the blank sample, and the Spectra Manager software 1.53 (JASCO) was used. Finally, CD spectra were plotted using GraphPad Prism 9 (GraphPad Software, Inc.).

### Size exclusion chromatography coupled with multi-angle light scattering (SEC-MALS)

Both recombinant proteins were analyzed by SEC-MALS after aggregates were removed by centrifugation. A Superdex 75 Increase 10/300 GL column (Cytiva) was firstly equilibrated with SEC buffer (50mM Tris, pH 7.0, 100mM NaCl). Then, the sample was applied to the column with a 0.5 mL/min flow rate on a 1260 Infinity II HPLC system (Agilent), which was coupled to a miniDAWN TREOS and Optilab T-rEX refractive index detector (Wyatt Technology). Protein samples were detected at 215 nm. Measurements were conducted in triplicates, and molecular mass distributions were calculated using the ASTRA v.7.3.0.18 software suite (Wyatt Technology).

### Ellman’s Assay

To prepare the Ellman’s reagent buffer, DTNB was dissolved in a buffer of 0.1M Na_3_PO_4_, 1mM EDTA, pH 8.0 at a final concentration of 4mg/ml. Then, cystine was dissolved in a buffer of 50mM Tris, pH 7.0, 100mM NaCl to a 1M concentration. This cystine solution was diluted to 200μM, 100μM, 50μM, 20μM, 10μM and 1μM. To generate the standard curve, 50μL of these solutions were added to 950μL of the Ellman’s reagent buffer. Protein samples were diluted to 40μM and 50μL of sample were added to 950μL of Ellman’s reagent buffer. For each recombinant protein, probes were measured in duplicate. All reactions were incubated at room temperature for 15min. Optical absorbance was measured at 412nm using an Ultrospec 2100 pro-operator (Amersham Biosciences). Plots were generated by GraphPad Prism 9 (GraphPad Software, Inc.).

