## Supplementary figures and images for "A unique Cysteine-type protein domain regulates cuticular extracellular matrix assembly in nematodes"

### Supplementary Figure 1

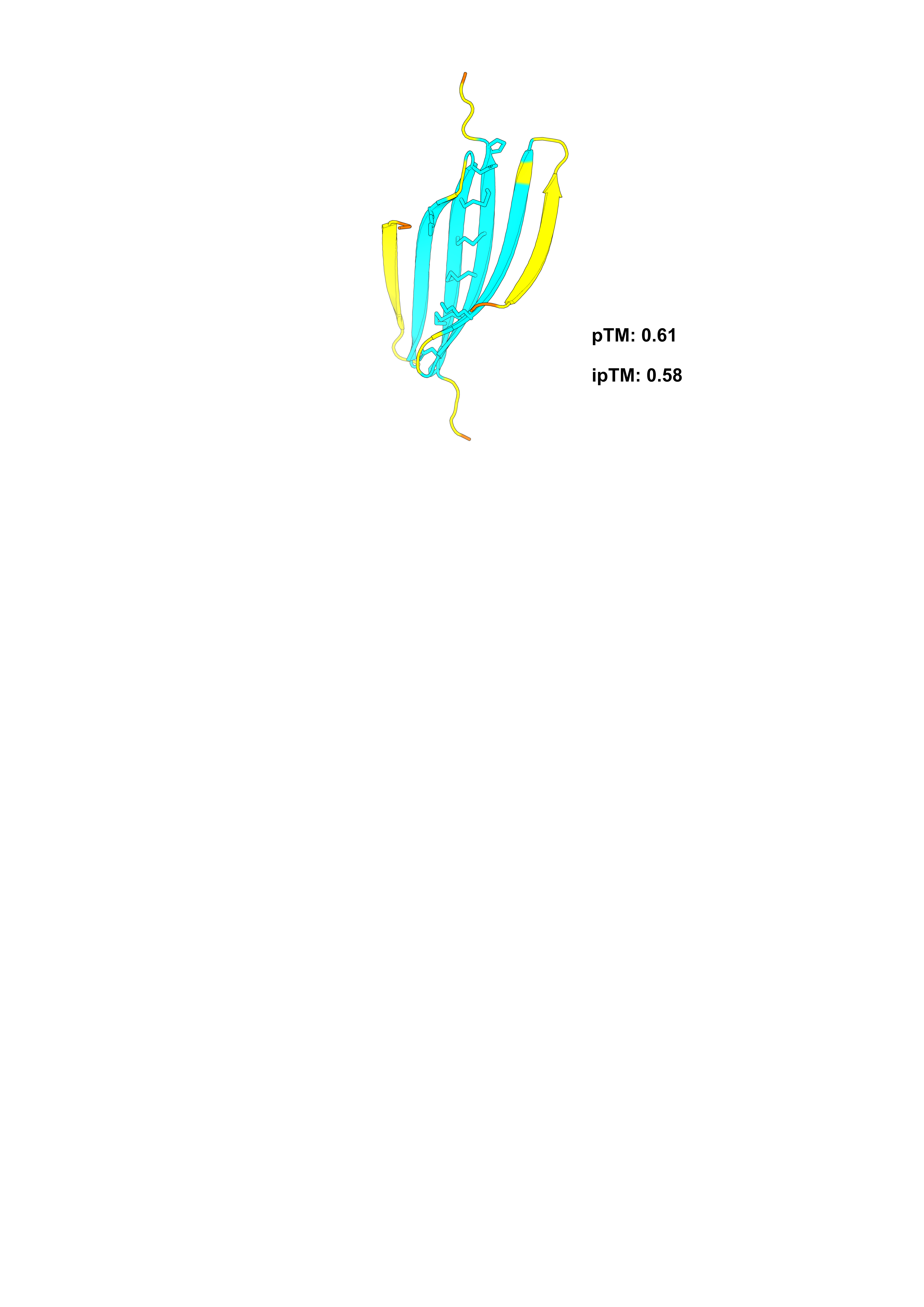

### Supplementary Figure 2

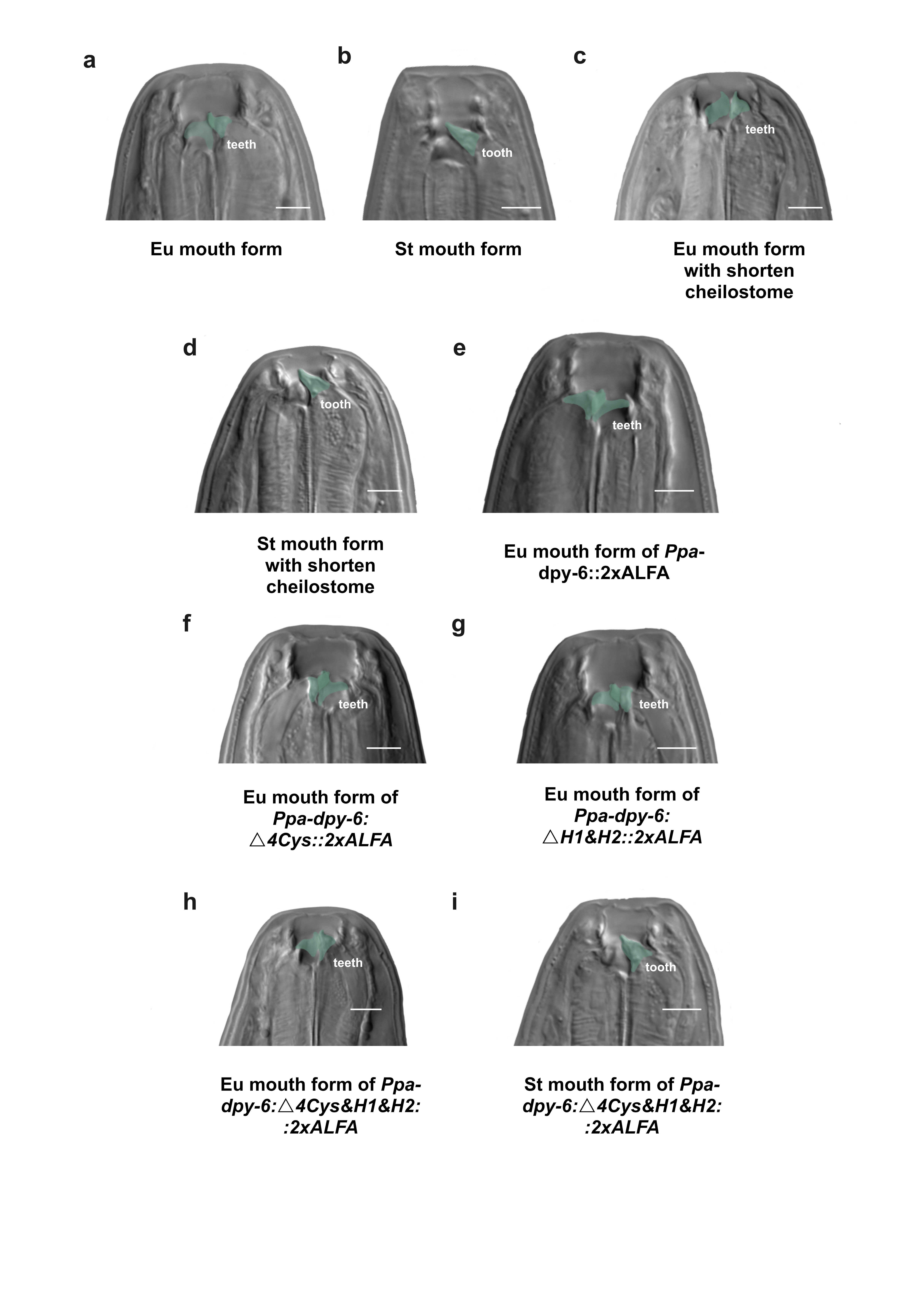
