## Supplementary Table 1&2&3 for "A unique Cysteine-type protein domain regulates cuticular extracellular matrix assembly in nematodes"

**Table 1. Amino acid boundary for each protein domain of DPY-6 in *P. pacificus* and *C. elegans***

|  | *P. pacificus* | *C. elegans* |
| --- | --- | --- |
| signal peptide | 1-20 | 1-20 |
| 4-Cys domain (disulfide staple domain) | 21-61 | 21-59 |
| coiled coil H1 | 195-228 | NA |
| coiled coil H2 | 256-287 | NA |
| long helical region | 1108-1203 | 1094-1195 |
| *Pristionchus*-specific insertion | 1204-1255 | NA |
| aECM cysteine cradle domain (aCCD) | 1256-1316 | 1196-1254 |

**Table. 2 *C. elegans* strains utilized in the study**

| Strain Name | tu number | RS number | Comments |
| --- | --- | --- | --- |
| N2 | NA | NA | wild type |
| *Cel*-dpy-6::2xALFA | tu2081 | RS4455 | ALFA-tagged full length DPY-6 protein |
| *Cel*-dpy-6:  △4Cys::2xALFA | tu2085 | RS4459 | In-frame deletion of disulfide staple domain |
| *Cel*-dpy-6 N-coil::  2X-ALFA | tu2259 | RS4789 | Introduction of coiled coils H1 and H2 from *P. pacificus* DPY-6 protein |
| *Cel*-dpy-6 N-coil  △4Cys::2X-ALFA | tu2271 | RS4803 | Introduction of coiled coils H1 and H2 from *P. pacificus*; in-frame deletion of disulfide staple domain |

**Table. 3 *P. pacificus* strains utilized in the study**

| Strain Name | tu number | RS number | Comments |
| --- | --- | --- | --- |
| PS312 | NA | NA | wild type |
| *Ppa*-dpy-6::2xALFA | tu2157 | RS4545 | ALFA-tagged full length DPY-6 protein |
| *Ppa*-dpy-6:  △4Cys::2xALFA | tu2185 | RS4580 | In-frame deletion of disulfide staple domain |
| *Ppa*-dpy-6:  △H1&H2::  2xALFA | tu2208 | RS4686 | In-frame deletion of coiled coils H1 and H2 |
| *Ppa*-dpy-6:△4Cys&  H1&H2::2xALFA | tu2232 | RS4757 | In-frame deletion of disulfide staple domain and coiled coils H1 and H2 |

**S-Figure 1. AlphaFold prediction of the P. pacificus disulfide staple domain**, showing pTM=0.61 and ipTM=0.58. Colorization follows pLDDT score (blue: pLDDT > 90; cyan: 90 > pLDDT > 70; yellow: 70 > pLDDT > 50; orange: 50 > pLDDT > 30).

**S-Figure 2.** **Mouth morphology of *P. pacificus* strains used in this study.** (a) Eurystomatous (Eu) mouth form of *P. pacificus*. (b) Stenostomatous (St) mouth form of *P. pacificus*. (c, d) Eu mouth form with a reduced cheilostome in both the Eu (c) and the St (d). (e) Eu mouth form of *Ppa*-dpy-6::2xALFA (tu2157). (f) Eu mouth form of *Ppa-dpy-6:△4Cys::2xALFA* (tu2185). (g) Eu mouth form of *Ppa*-dpy-6:△H1&H2::2xALFA (tu2208). (h) Eu mouth form of *Ppa*-dpy-6:△4Cys&H1&H2::2xALFA (tu2232). (i) St mouth form of *Ppa*-dpy-6:△4Cys&H1&H2::2xALFA (tu2232). Scale bars in all images are 5μm.
